# Somatosensory Perceptual Training Enhances Motor Learning by Observing

**DOI:** 10.1101/319905

**Authors:** Heather R. McGregor, Joshua G.A. Cashaback, Paul L. Gribble

## Abstract

Neuroimaging and neurophysiological studies in humans have demonstrated that action observation activates brain regions involved in sensory-motor control. A growing body of work has shown that action observation can also facilitate motor learning; observing a tutor undergoing motor learning results in functional plasticity within the motor system and gains in subsequent motor performance. However, the effects of observing motor learning extend beyond the motor domain. Converging evidence suggests that learning also results in somatosensory functional plasticity and somatosensory perceptual changes. This work has raised the possibility that the somatosensory system is also involved in motor learning that results from observation. Here we tested this hypothesis using a somatosensory perceptual training paradigm. If the somatosensory system is indeed involved in motor learning by observing, then improving subjects' somatosensory function before observation should enhance subsequent observation-related gains in motor performance. Subjects performed a proprioceptive discrimination task in which a robotic manipulandum moved the subject’s passive upper limb and he or she made judgments about the position of the hand. Subjects in a Trained Learning group received trial-by-trial feedback to improve their proprioceptive acuity. Subjects in an Untrained Learning group performed the same task without feedback. All subjects then observed a learning video showing a tutor adapting her reaches to a left force field (FF). We found that subjects in the Trained Learning group, who had superior proprioceptive acuity prior to observation, benefited more from observing learning compared to subjects in the Untrained Learning group. Improving somatosensory function can therefore enhance subsequent observation-related gains in motor learning. This study provides further evidence in favor of the involvement of the somatosensory system in motor learning by observing.

**Abbreviations:**

FF: :Force field
PD: :Maximum perpendicular deviation
IQR: :interquartile range

The authors report no financial interests or conflicts of interests.

## Introduction

Neuroimaging, neurophysiological, and behavioral studies provide evidence for a link between action observation and motor control in humans (Fadiga et al. 1995; Strafella and Paus 2000; Buccino et al. 2001; Flanagan et al. 2003; Watkins et al. 2003). A great deal of this work has focused on the potential role of an action-observation link in higher cognitive functions such as understanding and inferring the intentions of others’ actions (e.g., (Gallese and Goldman 1998; Rizzolatti et al. 2001). However, a growing body of research has also suggested a role for observation-related sensory-motor activity in motor learning (Heyes and Foster 2002; Mattar and Gribble 2005; Cross et al. 2006a; Alaerts et al. 2010; Buckingham et al. 2014).

A series of studies has demonstrated that action observation can facilitate force field (FF) adaptation (Mattar and Gribble 2005; Brown et al. 2009; Bernardi et al. 2013; Wanda et al. 2013). For this task, subjects grasp the handle at the end of a robotic manipulandum and adapt their reaching movements to forces applied by the robot (Shadmehr and Mussa-Ivaldi 1994). In a previous study, Mattar and Gribble (2005) presented subjects with a video showing another individual (“a tutor”) adapting his reaches to a robot-applied FF. Subjects who later performed reaches in the same FF as they had observed showed a benefit, performing straighter movements in the FF compared to non-observing subjects. Conversely, subjects who later performed reaches in the opposite FF to what they had observed showed a detriment, performing more curved movements in the FF compared to non-observing subjects. This study showed that subjects are able to learn about how to reach in novel FFs by observing a tutor’s movements (Mattar and Gribble 2005).

Motor learning by observing brings about functional changes in motor areas of the brain (Cross et al. 2009; Brown et al. 2009; McGregor and Gribble 2015; McGregor et al. 2016). However, the effects of observing motor learning are not restricted to the motor domain; there is accumulating evidence of observation-related neural and behavioral changes in the somatosensory domain as well (Bernardi et al. 2013; McGregor and Gribble 2015, 2017; McGregor et al. 2016; Valchev et al. 2017). For example, behavioral work has shown that observing motor learning alters somatosensory perception. Subjects performed a proprioceptive judgment task before and after observing a video of a tutor learning to reach in a FF. Observing motor learning not only facilitated subjects’ motor performance in the observed FF, but it also altered subjects’ proprioceptive judgments. Observing motor learning resulted in small but systematic changes in subjects’ perceived limb position depending on the FF that had been observed (Bernardi et al. 2013). These results suggested that observing motor learning affects not only the motor system, but also the somatosensory system.

Our recent work is consistent with the idea that the somatosensory system plays a role in motor learning by observing. Using EEG, we measured changes in primary somatosensory cortex (S1) excitability after observing motor learning. We found that somatosensory evoked potentials increased from pre- to post-observation and did so for those subjects who had observed a tutor adapting to a FF, but not for subjects who had observed similar movements that did not involve learning. Furthermore, post-observation increases in S1 excitability were correlated with subsequent behavioral measures of motor learning. These results suggest that observation-induced functional changes in S1 are involved in motor learning by observing (McGregor et al. 2016).

In a follow-up experiment, we showed that interfering with somatosensory cortical processing during observation can disrupt observation-related gains in motor learning. We applied electrical stimulation to the median nerve while subjects observed a video showing a tutor undergoing FF learning. The purpose of this experimental manipulation was to occupy the somatosensory system with afferent inputs that were unrelated to the observed learning task. During observation, subjects received median nerve stimulation to either the right arm (the same arm used by the tutor in the video), to the left arm, or no stimulation. Stimulation of the right arm (the observed effector) interfered with learning whereas stimulation applied to the left arm did not (McGregor et al. 2016). These findings are consistent with the idea that a somatosensory representation of the observed effector plays an important role in motor learning by observing; it must be available and unoccupied during observation in order for learning to be achieved.

If the somatosensory system is indeed involved in motor learning by observing, as the studies above suggest, then we predicted that improving subjects’ somatosensory function prior to observation should enhance subsequent motor learning by observing. We tested this idea in the current study by using a perceptual training paradigm to improve subjects’ sense of limb position prior to observation. Subjects performed a proprioceptive discrimination task in which a robotic arm displaced the hand and subjects made judgments about the relative location of the hand in the absence of visual feedback. Subjects in a Trained group received trial-by-trial feedback about the accuracy of limb position judgments during the proprioceptive task. Trial-by-trial feedback was withheld from an Untrained group. Subjects then observed a video showing a tutor adapting to a FF. Finally, subjects performed reaches in a FF as a behavioral assessment of motor learning by observing. We found that providing trial-by-trial feedback during the proprioceptive discrimination task increased subjects’ proprioceptive acuity. The post-observation behavioral assessment revealed that the Trained group, who had superior proprioceptive acuity prior to observing motor learning, benefited more from observation compared to the Untrained group. This finding is consistent with the idea that somatosensory perceptual training improves proprioceptive function which in turn enhances motor learning by observing.

## Methods

### Subjects

Seventy-eight subjects participated in this experiment. Subjects were assigned to one of three groups: a Trained Learning group (n = 26, 8 males, mean age = 21.6 years ± 0.65 years SEM), an Untrained Learning group (n = 26, 9 males, mean age = 21.4 ± 0.58 years SEM) or a Trained Control group (n = 26, 8 males, mean age = 20.8 years ± 0.44 years SEM). All subjects were right handed, had normal or corrected-to-normal vision, and were naïve to robot-imposed force fields. Subjects reported no neurological or musculoskeletal disorders. Subjects provided written informed consent to experimental procedures approved by the Research Ethics Board at the University of Western Ontario.

### Apparatus

Subjects were seated in front of a custom tabletop and grasped the handle of a two-joint, two degree-of-freedom robotic arm (InMotion2, Interactive Motion Technologies) with their right hand. The right arm was abducted approximately 80° from the trunk and was secured atop an air sled, which supported the arm against gravity. A liquid crystal display (LCD) TV projected visual feedback onto a semi-silvered mirror mounted horizontally above the robotic arm during the reaching task.

### Reaching Task

During the reaching task, subjects were instructed to guide the robot handle forward in a straight line from a home position (20-mm blue circle) to a single visual target (20-mm white circle). The position of the robot handle was represented by a 5-mm pink circular cursor. Upon the completion of each reach, the target changed color to provide subjects with feedback about movement timing. The target disappeared if the movement was completed with the desired time (375 ± 100 ms). The target turned red or green to indicate that duration of a movement was too slow or too long, respectively. Following movement timing feedback, the robot moved the subject’s passive arm to the home position to begin the next trial.

The robotic arm applied a velocity-dependent force field (FF) during the reaching task according to the following equation:

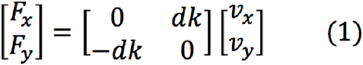

in which x and y are lateral and sagittal directions, Fx and Fy are the robot forces applied at the hand, v_x_ and v_y_ are hand velocities, k=14 Ns/m, and d=+1 (right FF), −1 (left FF) or 0 (null field).

### Reaching Video Stimuli

Two videos were used in the study, each showing a tutor performing the reaching task described above from a top-down perspective (McGregor and Gribble 2015). A learning video consisted of a series of 30-second clips showing a tutor adapting her reaches to a left FF. This video depicted highly curved movements that gradually straightened as the tutor adapted to the FF. A control video consisted of a series of 30-second clips showing a tutor performing reaches in an unlearnable randomly-varying FF. The direction of the force field in this video varied randomly from trial-to-trial between a left FF, right FF, or a null field. The control video therefore showed both high and low curvature movements, but lacked the progressive decrease in movement curvature that was depicted in the learning video. Each video showed a total of 200 reaches and was 15 minutes in duration (including regular breaks).

### Proprioceptive Discrimination Task

We used a proprioceptive discrimination task to assess subjects’ proprioceptive acuity (sensitivity to displacements in limb position). We used a two-alternative forced choice task in which subjects made judgments about the relative position of their hand (Ostry et al. 2010; Wong et al. 2011). Subjects held onto the handle of the robotic arm with their right hand, closed their eyes and relaxed their right arm, which was supported by an air sled. Each trial consisted of a reference phase, a passive movement phase, and a judgment phase.

During the reference phase, the robot held the subject’s hand at the central reference position for 2 seconds. Next, in the passive movement phase, the robot moved the subject’s hand away from the reference and back (along a medial-lateral axis), stopping at a test location near the reference. Subjects were instructed not to resist movement generated by the robot. The aim of the passive movement phase was to bring the subject’s hand from the central reference position to the test location without providing cues that could be used in aiding subject’s subsequent judgments. Features of the passive movement were randomized from trial to trial including: movement direction (left or right), total path length (14 ± 2 cm SD), and movement duration (1–1.6 s). Movements of the passive limb by the robot followed a minimum jerk trajectory (Flash and Hogan 1985).

During the hand position judgment phase, the robot held the subject’s hand at the test location and the subject reported whether his or her hand was located to the left or to the right of the reference position. Following the subject’s response, the robot moved the hand back to the reference position via an indirect path (along a left-right axis). The direction, path length, and duration were randomized such that subjects were not provided with cues about the accuracy of their judgment on the previous trial.

Each block of the proprioceptive discrimination task consisted of 74 trials. We presented test locations at 7 distances from the reference position (0 ± 0.67, 1.33, and 3.0 cm). Test locations were presented using the method of constant stimuli with the following frequencies: 0 cm (14 trials), ± 0.67 cm (12 trials each), ± 1.33 cm (12 trials each), and ± 3 cm (6 trials each). The ± 3 cm test locations were presented less frequently because subjects typically respond with a 100% judgment accuracy at these test locations.

All subjects performed 5 blocks of the proprioceptive discrimination task (370 proprioceptive judgment trials in total). Subjects were given breaks halfway through each block and between blocks (i.e., every 37 trials). During each break, all subjects were told their percent accuracy over the previous 37 trials. To motivate subjects throughout the proprioceptive task blocks, we offered a performance-based monetary bonus of up to $10 CAD in addition to the hourly base rate of compensation.

### Experimental Design

Each subject participated in one 2-hour session (see Figure 1A). The experimental session began with subjects performing 30 practice reaches in a null field (no force applied, data not shown). Subjects then performed 50 reaches to the visual target in the null field. This allowed us to assess subjects’ baseline movement curvature.

**Figure 1:**
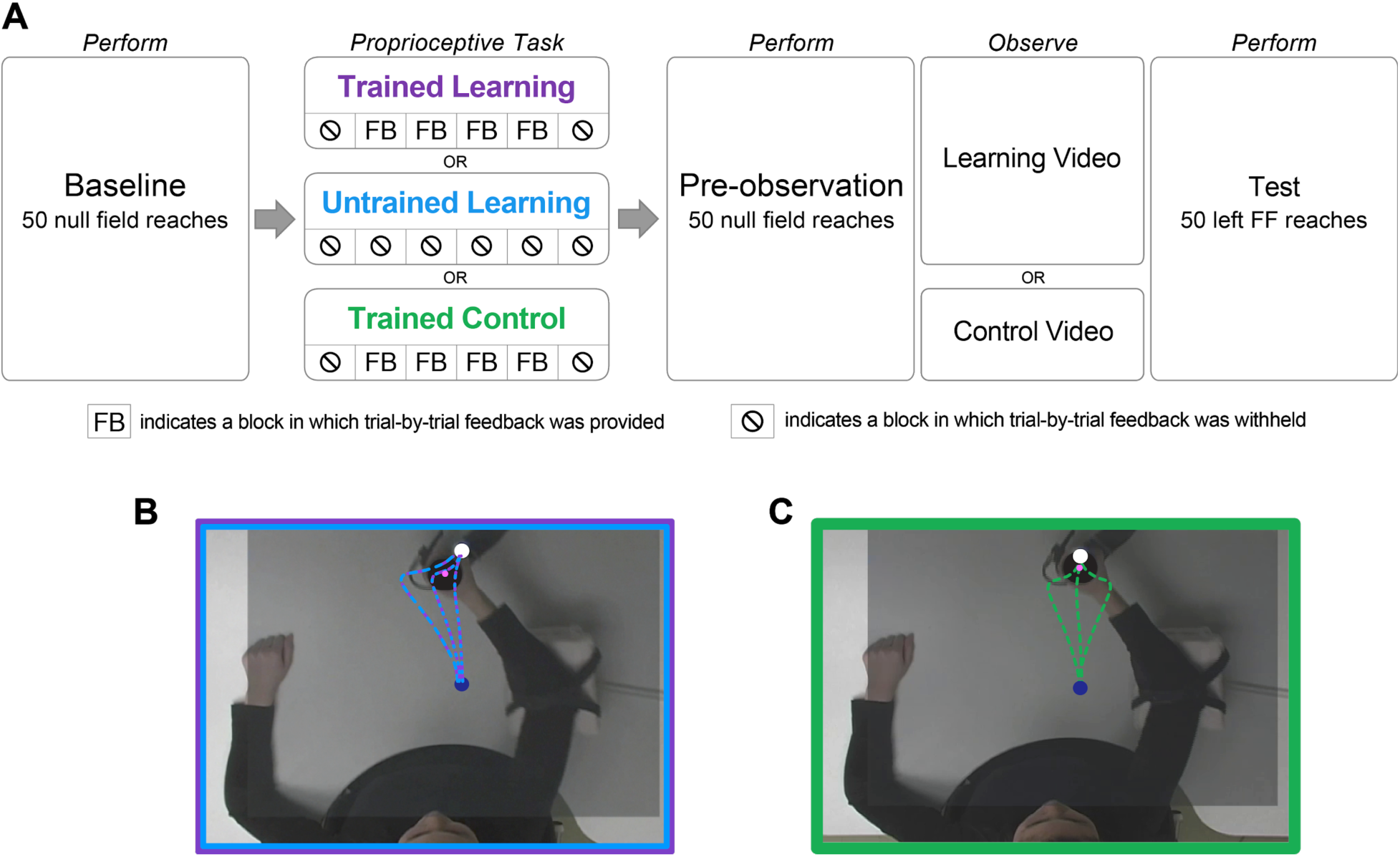
A, Experimental design. Subjects grasped the handle of the robotic arm with their right hand. First, subjects performed 50 reaches in a null field (no force applied by the robot). Next, all subjects performed the proprioceptive discrimination task in which the robot moved the passive limb and subjects judged whether their hand was displaced to the left or to the right of a central reference position. During blocks 1 and 5, trial-by-trial judgment accuracy feedback was withheld from all subjects. During blocks 2, 3 and 4, subjects in the Trained Learning group and the Trained Control group received proprioceptive training. For both of the Trained groups, the experimenter provided verbal accuracy feedback immediately following each reported judgment, informing the subject if the judgment was correct or incorrect (and if incorrect, providing the correct answer). The Untrained Learning group did not receive trial-by-trial feedback during any of the blocks. All subjects then performed 50 null field reaches. Subjects in the Trained Learning and Untrained Learning groups observed a learning video showing a tutor adapting her reaches to a left force field (FF). Subjects in the Trained Control group observed a control video showing a tutor performing reaches in an unlearnable randomly-varying FF. Finally, all subjects performed reaches in a left FF as a behavioral assessment of motor learning by observing. **B, Screenshot of the learning video.** The learning video showed a tutor adapting her reaches to a left FF. Superimposed hand trajectories are for demonstrative purposes only and were not shown in the experiment. **C, Screenshot of the control video.** The control video showed a tutor performing reaches in an unlearnable FF in which the direction of the applied force varied randomly from trial-to-trial. Superimposed trajectories are for demonstrative purposes only.

Subjects were then assigned to one of three groups, two of which received perceptual training (‘Trained’ groups) and one did not (‘Untrained’ group). All subjects performed 5 blocks of the proprioceptive discrimination task described above. During blocks 1 and 5, trial-by-trial accuracy feedback was withheld from all subjects. During blocks 2–4, Trained subjects received trial-by-trial feedback. Immediately after making a verbal response (“left” or “right”), the experimenter informed the subject if the judgment was correct or if it was incorrect, and then informed him or her of the correct response. During blocks 2–4, subjects in an Untrained group continued performing the proprioceptive task without trial-by-trial accuracy feedback. The idea was to improve proprioceptive acuity of subjects in both of the Trained groups, but not for subjects in the Untrained group. Regardless of their training condition, all subjects were told their percent accuracy during each break (i.e., every 37 trials).

Following the proprioceptive discrimination task, all subjects performed a second set of 50 reaches in the null field. This allowed us to test whether the perceptual training itself resulted in changes in movement curvature.

Next, the Untrained group and one group of Trained subjects observed the learning video (Untrained Learning group and Trained Learning group, respectively) which showed a tutor adapting her reaches to a left FF. The remaining group of Trained subjects observed the control video (‘Trained Control’ group) which showed a tutor performing reaches in an unlearnable randomly-varying FF. During observation, all subjects remained still with their arms rested on the tabletop beneath the robot arm. To ensure that subjects paid attention to the video, they were instructed to count the number of times the tutor in the video performed a reach within the desired time range (indicated by the target disappearing). Subjects reported their tallies during video breaks. Reported tallies were not incorporated into data analyses. Reported tallies were over 95% accurate on average for all groups, and subjects were not excluded based on their reported tallies.

Finally, we assessed motor learning by observing by having all subjects perform reaches in a left FF. The more subjects learned about the left FF from observing the learning video, the better, straighter reaches they would perform when they later encounter that same left FF. Therefore, lower movement curvature in the left FF would indicate greater motor learning by observing.

### Proprioceptive Data Analysis

Here we assessed if improving proprioceptive function prior to observation could enhance motor learning by observing. We estimated each subject's proprioceptive acuity on the basis of his or her binary judgment data from blocks 1 and 5 of the proprioceptive discrimination task, during which trial-by-trial feedback was withheld from all subjects. For each of these blocks, we estimated a sigmoidal psychometric function based on the subject’s binary response data. Proprioceptive acuity was quantified as the interquartile range (IQR) of the psychometric function, a measure also known as uncertainty range (Henriques and Soechting 2003). This measure of uncertainty is inversely related to acuity such that a smaller IQR value indicates that the slope of the psychometric function is steep and therefore the subject is sensitive to small displacements in limb position. We predicted that providing trial-by-trial feedback during blocks 2–4 of the proprioceptive discrimination task would improve perceptual acuity (and hence decrease IQR values) from block 1 to 5 for the Trained Learning and Trained Control groups.

Changes in proprioceptive acuity were assessed using a split-plot analysis of variance (ANOVA) followed by planned pairwise comparisons. The dependent measure for the ANOVA was the IQR of the psychometric function. The ANOVA used group (Trained Learning, Untrained Learning, Trained Control) as the between-subjects factor and proprioceptive task block (1, 5) as the within-subject factor. We also examined changes in subjects’ judgment accuracy from block 1 and block 5. For this analysis, we performed a split-plot ANOVA using group (Trained Learning, Untrained Learning, Trained Control) as the between-subjects factor and proprioceptive task block (1, 5) as the within-subject factor. The dependent measure was the percentage of correct judgments within each of the blocks.

### Motor Behavior Analysis

Robot handle positions were sampled at 600 Hz. Velocities were computed using a central difference algorithm. Positions and velocities were low-pass filtered offline using a second order Butterworth filter implemented in MATLAB (Mathworks, Inc.) with a cutoff frequency of 40 Hz. For each trial, we computed the maximum point of lateral deviation of the hand path relative to a straight line connecting the home and target. This measure is known as the maximum perpendicular deviation (PD). We then computed a motor learning by observing score for each subject. This measure was computed as the average PD of the subject’s first 3 reaches in the left FF minus the average PD of the last 25 reaches in the baseline null field condition. This measure therefore reflects the extent to which the subject’s performance in the left FF benefited from observation relative to his or her baseline PD in the null field. As we have demonstrated previously (Mattar and Gribble 2005; Brown et al. 2009), learning about a FF from observation results in better, straighter movements when subjects later perform reaches in the same FF. As such, we predicted that greater motor learning by observing would result in straighter movements in the left FF and therefore higher (i.e., closer to zero) motor learning by observing scores. Group differences in motor learning by observing scores were assessed using a one-way between-subjects ANOVA.

## Results

### Proprioceptive Training Results

We tested for changes in IQR using a split-plot ANOVA, which revealed a group x proprioceptive test block interaction (F(2,75) = 3.72, p = 0.03; Figure 2A). Subjects in the Trained Learning group and the Trained Control group were exposed to the same experimental protocols by the end of the proprioceptive discrimination task. The only difference between the protocol used for these two groups was the video that was observed *following* proprioceptive training. Therefore, the observed IQR changes depended on whether or not proprioceptive judgment feedback had been provided during blocks 2 through 4. The Trained Learning group exhibited greater IQR decreases compared to subjects in the Untrained Learning group (t(50) = - 2.47, p = 0.008). Similarly, the Trained Control group exhibited greater IQR decreases compared to subjects in the Untrained Learning group (t(50) = −2.25, p = 0.02). No reliable differences in post-training IQR decreases were observed between the Trained Learning and Trained Control groups (t(50) = 0.06, p = 0.95). Changes in IQR are further illustrated in Figure 2B which shows the average psychometric fit for each group during blocks 1 and 5. The IQR of a psychometric function is inversely related to perceptual acuity, with smaller IQR values indicating greater sensitivity to displacements in limb position. Prior to observation, subjects in both of the Trained groups therefore exhibited superior proprioceptive acuity compared to subjects in the Untrained group.

**Figure 2:**
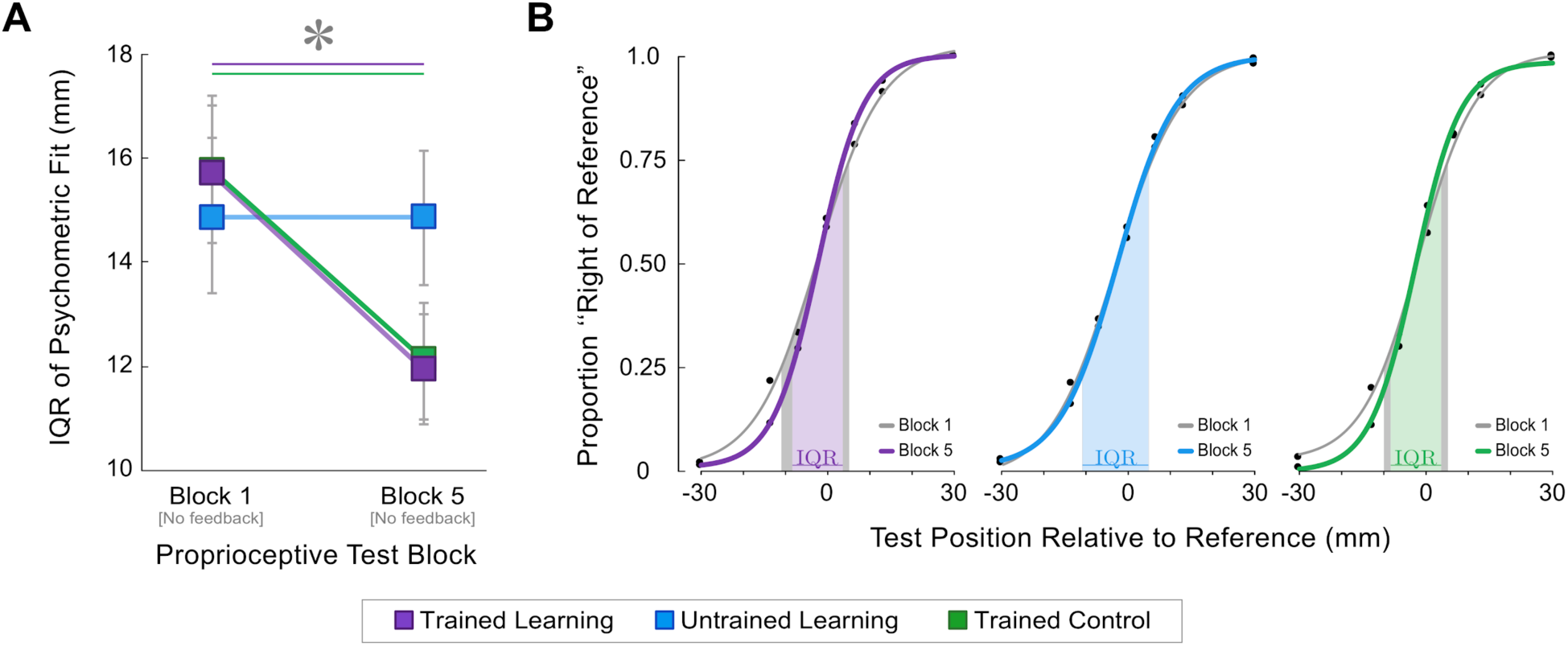
Proprioceptive training increased proprioceptive acuity. **A**, Perceptual acuity was assessed in blocks 1 and 5 of the proprioceptive discrimination task, during which trial-by-trial feedback was withheld from all subjects. Perceptual acuity was quantified as the interquartile range (IQR) of the psychometric function estimated from each subject’s set of binary responses. In block 5, only the Trained Learning group (purple) and the Trained Control group (green) exhibited reliable IQR decreases. Error bars represent SE. * indicates p < 0.05. **B**, Group average psychometric fits for blocks 1 and 5 of the proprioceptive discrimination task. The IQR of each fit is indicated by the shaded areas along the x-axis. IQR decreased from block 1 to block 5 only for the Trained Learning group (left, purple) and the Trained Control group (right, green).

### Motor Behavior Results

Training subjects on the proprioceptive discrimination task resulted in improvements to perceptual acuity. We tested if trained subjects with superior somatosensory performance prior to observation go on to achieve greater motor learning by observing. Following the proprioceptive task, subjects observed either a learning video showing a tutor adapting to a left FF or a control video showing a tutor reaching in an unlearnable FF. Following observation, we assessed the extent to which subjects learned from observation by instructing them to perform reaches in a left FF (the same FF that was shown in the learning video). As in previous work (Mattar and Gribble 2005; Brown et al. 2009; Bernardi et al. 2013), we expected that motor learning by observing would primarily affect initial performance in the left FF, after which all groups would adapt to the left FF through physical practice.

Figure 3A shows average learning curves in the left FF for each group. It can be seen that the Trained Learning group, who had superior proprioceptive acuity prior to observation, performed straighter movements when first exposed to the left FF compared to subjects in the Untrained Learning group. This is consistent with the idea that superior proprioceptive acuity prior to observation enhanced the extent to which subjects benefited from observing learning. However, it is possible that the Trained Learning group’s straighter movements in the left FF were due to general increases in movement straightness following perceptual training (and not to motor learning by observing). Superior proprioceptive acuity may have made subjects in the Trained Learning group more sensitive to felt displacements in limb position during left FF reaches and allowed for faster movement corrections. Therefore, it is feasible that increased perceptual acuity alone could account for the observed group differences in left FF performance. If this were the case, then we would predict that increasing proprioceptive acuity would result in similar movement curvature in the left FF as that of the Trained Learning group regardless of the video that was observed. We tested this idea by running a Trained Control group. Subjects in the Trained Control group showed similar post-training increases in proprioceptive acuity as the Trained Learning group following the proprioceptive discrimination task. However, after observing the control video, subjects in the Trained Control group performed movements in the left FF that were more curved than either the Trained Learning group or Untrained Learning group (Figure 3A).

**Figure 3:**
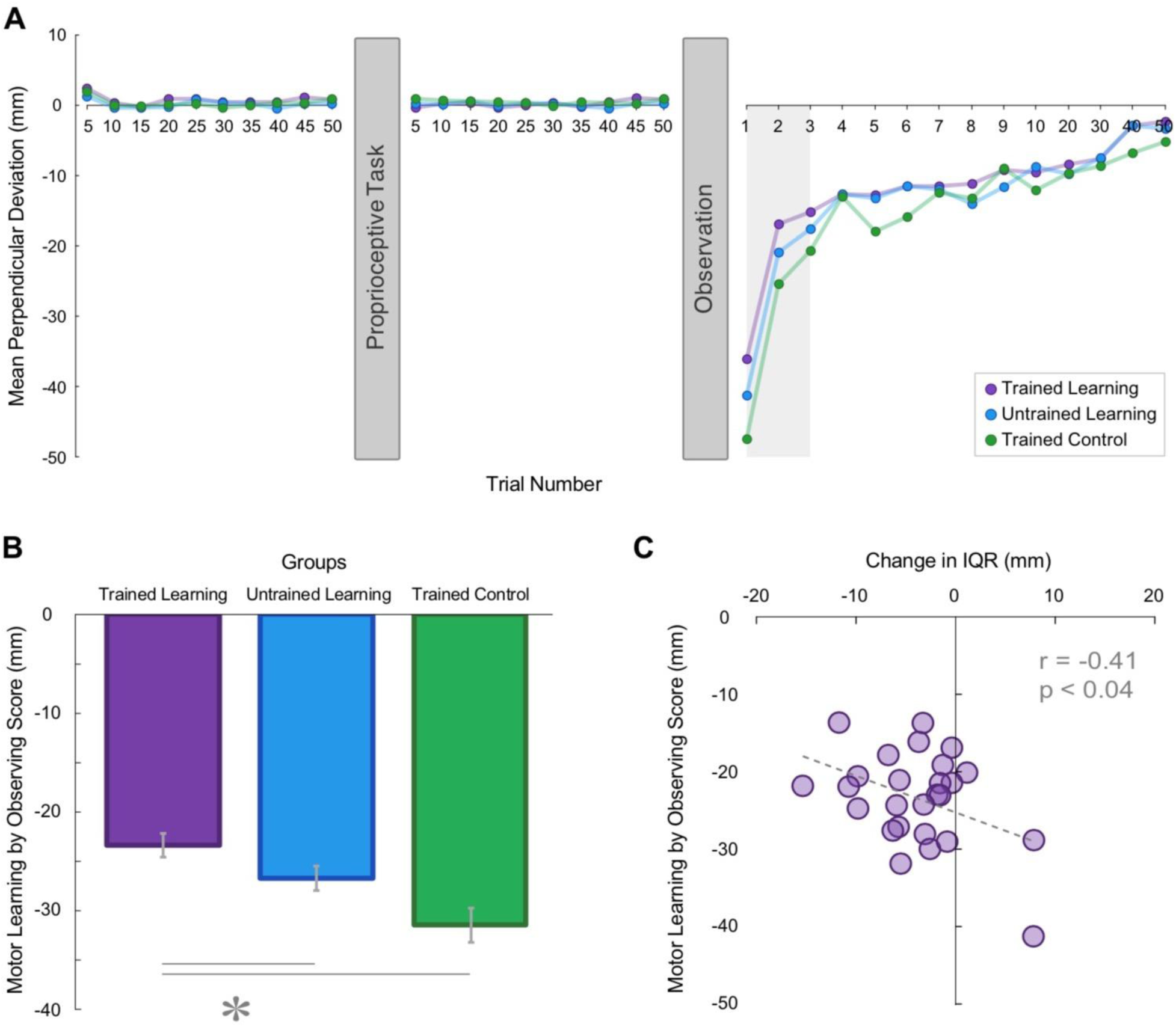
Proprioceptive training enhanced subsequent motor learning by observing. **A**, Evolution of perpendicular deviation (PD). The first 10 data points correspond to individual trial group averages. Data points thereafter correspond to the group averages of 5-trial blocks. The grey shaded region of the plot indicates the first 3 trials in the left FF, which were used to calculate motor learning by observing scores. Those subjects who observed the learning video (i.e., Trained Learning and Untrained Learning groups) showed a benefit, performing straighter reaches in the same (left) FF. **B**, Motor learning by observing scores as a function of video observed. Motor learning by observing scores reflect perpendicular deviation of initial movements in the left FF. Subjects in the Trained Learning and Untrained Learning groups therefore exhibited higher motor learning by observing scores (i.e., they performed less curved movements in the left FF) compared to the Trained Control group. Moreover, subjects in the Trained Learning group exhibited higher motor learning by observing scores than the Untrained Learning group. * indicates p < 0.05. Error bars represent SE. **C**, Across subjects in the Trained Learning group, post-training decreases in the IQR of subjects’ psychometric fit correlated with subsequent behavioral measures of motor learning by observing. Post-training increases in perceptual acuity corresponded to the extent to which subjects subsequently benefited from observing the left FF.

We quantified performance in the left FF by computing motor learning by observing scores (Figure 3B). A one-way between-subjects ANOVA revealed reliable group differences in motor learning by observing scores (F(2,75) = 8.39, p = 0.01). The Trained Learning group exhibited higher motor learning by observing scores compared to the Untrained Learning group (t(50) = 1.94, p = 0.028), even though both groups had observed the same learning video. Moreover, the movements of the Trained Control group were more curved in the left FF compared to those of either the Trained Learning group (t(50) = 3.87, p < 0.001) or the Untrained Learning group (t(50) = 2.24, p = 0.015).

We examined if proprioceptive acuity at the end of perceptual training (prior to observation) was correlated with subsequent motor learning by observing. However, there was no reliable correlation between post-training IQR values and subsequent behavioural motor learning by observing scores for either the Trained Learning group (r = −0.22, p = 0.28), the Untrained Learning group (r = −0.06, p = 0.78), or the Trained Control group (r = −0.25, p = 0.23). Similarly, there was no reliable correlation between pre-training IQR values and subsequent behavioural scores of motor learning by observing for either the Trained Learning group (r = 0.147, p = 0.47), the Untrained Learning group (r = −0.070, p = 0.73), or the Trained Control group (r = −0.403, p = 0.245).

We further examined if the extent to which proprioceptive acuity changed following the proprioceptive task and motor learning by observing. As shown in Figure 3C, across subjects in the Trained Learning group, there was a statistically reliable correlation between post-training changes in the IQR of the psychometric fit and subsequent behavioral measures of motor learning by observing (r = −0.41, p = 0.036). That is, those subjects who showed the greatest improvements in perceptual acuity following perceptual training were those who went on to benefit more from observing motor learning. No reliable relationship between perceptual acuity change and observation-related changes in movement was seen for subjects in the Untrained Learning group (r = 0.004, p = 0.98) and the Trained Control group (r = 0.25, p = 0.22).

## Discussion

We tested the idea that somatosensory perceptual training improves somatosensory function and this in turn enhances subsequent motor learning by observing. Subjects underwent perceptual training on a proprioceptive discrimination task to improve their proprioceptive acuity. A Trained Learning group who, on average, possessed superior proprioceptive acuity before observation benefited more from observing motor learning compared to subjects to subjects in an Untrained Learning group. For subjects in the Trained Learning group, post-training increases in proprioceptive acuity were reliably correlated with subsequent behavioral measures of motor learning by observing.

The Trained Learning group’s superior performance (i.e., straighter movements) in the left FF was not due to observing motor learning alone or due to perceptual training alone. If that were the case, we would have found comparable motor performance in the left FF across all three groups. Rather, our results show that the Trained Learning group’s performance benefit was due to the combination of perceptual training *and* observing motor learning. This suggests that improving proprioceptive acuity prior to observation can enhance subsequent observation-related gains in motor learning.

The idea that the somatosensory system is involved in motor learning by observing is supported by previous behavioral work demonstrating that observing motor learning alters somatosensory perception. Bernardi and colleagues (2013) examined proprioceptive function before and after participants observed a video of a tutor learning to reach in a FF. Proprioceptive function was assessed using a discrimination task in which a robot manipulandum moved the hand away from the body along one of several trajectories and the subject judged whether the hand had been displaced to the left or the right (in the absence of visual feedback). Bernardi and colleagues reported that observing motor learning not only facilitated subjects’ motor performance in the observed FF, but it also altered subjects’ judgments of perceived limb position. Observing motor learning resulted in systematic changes in subjects’ somatosensory perception depending on the FF that had been observed. Observing a video depicting right FF learning changed subjects’ proprioceptive perception such that judgments were biased toward the right. Conversely, observing a video depicting left FF resulted in proprioceptive judgments being biased toward the left. These results suggested that observing motor learning affects not only the motor system, but also the somatosensory system.

Using resting-state fMRI, we have shown that observing FF learning indeed results in functional changes to the somatosensory brain areas in addition to visual and motor areas of the brain (McGregor and Gribble, 2015). We assessed changes in resting-state functional connectivity from pre- to post- observation that were related to behavioral measures of motor learning by observing. Observing motor learning changed functional connectivity between primary somatosensory cortex, visual area V5/MT, the cerebellum, and primary motor cortex. Observation-induced functional connectivity changes within this network were correlated with subsequent behavioral measures of motor learning by observing. That is, those subjects who showed greater functional connectivity changes after observing learning were those who achieved greater motor learning from observation (McGregor and Gribble 2015).

Vahdat and colleagues (2014) have recently showed that a perceptual training protocol similar to the task used in the current study increases resting-state functional connectivity among somatosensory, motor, and premotor brain areas. In their variation of the perceptual training task, a robotic manipulandum moved the subject’s hand away from the body along one one of several fan-shaped trajectories and the subject judged whether the hand was displaced to the left or to the right of the midline. As in the current study, providing reinforcement feedback during this task resulted in improvements to acuity of sensed limb position. Resting-state fMRI data acquired before and after perceptual training showed increases in functional connectivity between bilateral S1, left M1, dorsal premotor cortex, and the superior parietal lobule which correlated with behavioural measures of post-training perceptual improvements (Vahdat et al. 2014). This network bears a strong resemblance to the network we have previously reported in which pre-observation functional connectivity predicts subsequent motor learning by observing (McGregor and Gribble, 2017). Given the similarity between the perceptual training protocol used by Vahdat et al (2014) and the protocol used the current study, it is likely that our perceptual training induced functional changes within a similar network of somatosensory, motor and premotor brain areas. Our previous work has suggested that a somatosensory representation associated with the observed effector (i.e., right arm) plays a necessary role in motor learning by observing (McGregor et al. 2016). Taken together, it is possible that somatosensory perceptual training primed somatosensory, motor, and premotor areas for subsequent observational learning.

More generally, the results of the current study complement the findings of studies of motor learning, in which subjects learn by performing a task through physical practice. There is accumulating evidence that motor learning involving physical practice results in changes to somatosensory perception. Motor learning can result in perceptual acuity improvements (Wong et al. 2011) as well as changes in sensed limb position (Cressman and Henriques 2009; Ostry et al. 2010; Haith et al. 2009). For example, FF adaptation has been shown to change sensed limb position, shifting it left or right depending on the direction of the learned FF (Ostry et al. 2010).

Neuroimaging and EEG studies further complement the evidence that functional changes in somatosensory brain areas occur with motor learning (Vahdat et al. 2011; Nasir et al. 2013). Using resting-state fMRI, Vahdat and colleagues (2011) showed that motor learning alters functional connectivity involving somatosensory brain areas. Undergoing FF learning increased functional connectivity between second somatosensory cortex, ventral premotor cortex, and supplementary motor area. Moreover, the degree to which functional connectivity increased was correlated with behavioral measures of learning-related changes to sensed limb position.

Somatosensory perceptual training has also been shown to enhance subsequent motor learning through physical practice (Rosenkranz and Rothwell 2012; Wong et al. 2012; Darainy et al. 2013; Vahdat et al. 2014). Darainy and colleagues (2013) trained subjects on a proprioceptive task similar to the one used in the present study, in which a robotic manipulandum moved the hand from a reference along one of several fan-shaped trajectories and subjects made judgments regarding the displacement of the hand. When provided with reinforcement accuracy feedback during the proprioceptive task, subjects showed increased perceptual acuity as well as decreases in perceptual bias such that they were more accurate in perceiving the boundary between left and right. They also found that perceptual training resulted in improvements in subsequent FF adaptation, improving the rate of learning, the extent of learning, and measures of predictive compensatory forces (assessed based on subject’s applied forces during error clamp trials). As we found in the current study, post-training increases in proprioceptive acuity were correlated with improvements in motor learning (Darainy et al. 2013).

While much of the work on motor adaptation has focused on the roles of M1 and the cerebellum, a recent optogenetic study has suggested a causal role for S1. Mathis and colleagues (2017) used a modified version of a FF reaching task in which mice grasped a joystick and adapted their pulling movements to applied forces. To test the role of S1 in FF adaptation, the authors photoinhibited the forelimb area of contralateral S1 partway through an adaptation block. S1 inhibition had no effect on previously adapted movements, but prevented mice from adapting further. The authors then showed that the detrimental effects of S1 inhibition were specific to motor adaptation. S1 photoinhibition did not impair reaches performed in the absence of applied forces, and mice were still able to learn a reinforcement-based task. These findings suggest that S1 plays a critical role in motor learning (Mathis et al. 2017).

Here we showed that improving subjects’ proprioceptive acuity through perceptual training prior to observation increased observation-related gains in motor learning. Moreover, post-training increases in proprioceptive acuity were correlated with subsequent behavioral measures of motor learning. Improving somatosensory function (i.e., proprioceptive acuity) can therefore enhance motor learning through observation. Somatosensory perceptual training may prime the sensory-motor system, thereby facilitating subsequent observational learning. The findings of this study further support the idea that the somatosensory system plays a role in motor learning by observing.

## Acknowledgements

This work was supported by the Natural Sciences and Engineering Research Council of Canada, the Canadian Institutes for Health Research, and by the National Institute of Child Health and Human Development R01 HD075740.

